# Harmonizing semantic annotations for computational models in biology

**DOI:** 10.1101/246470

**Authors:** ML Neal, M König, D Nickerson, G Mısırlı, R Kalbasi, A Dräger, K Atalag, V Chelliah, M Cooling, DL Cook, S Crook, M de Alba, SH Friedman, A Garny, JH Gennari, P Gleeson, M Golebiewski, M Hucka, N Juty, N Le Novère, C Myers, BG Olivier, HM Sauro, M Scharm, JL Snoep, V Touré, A Wipat, O Wolkenhauer, D Waltemath

**Author notes:** Corresponding author: Maxwell Lewis Neal, Center for Infectious Disease Research, 307 Westlake Ave. N, Suite 500, Seattle, WA 98109.

## Abstract

Life science researchers use computational models to articulate and test hypotheses about the behavior of biological systems. Semantic annotation is a critical component for enhancing the interoperability and reusability of such models as well as for the integration of the data needed for model parameterization and validation. Encoded as machine-readable links to knowledge resource terms, semantic annotations describe the computational or biological meaning of what models and data represent. These annotations help researchers find and repurpose models, accelerate model composition, and enable knowledge integration across model repositories and experimental data stores. However, realizing the potential benefits of semantic annotation requires the development of model annotation standards that adhere to a community-based annotation protocol. Without such standards, tool developers must account for a variety of annotation formats and approaches, a situation that can become prohibitively cumbersome and which can defeat the purpose of linking model elements to controlled knowledge resource terms. Currently, no consensus protocol for semantic annotation exists among the larger biological modeling community. Here, we report on the landscape of current semantic annotation practices among the COmputational Modeling in BIology NEtwork (COMBINE) community and provide a set of recommendations for building a consensus approach to semantic annotation.

## Introduction

Biological researchers use computational models to articulate hypotheses about the organization and dynamics of biological systems. Modeling has become an essential component of life science research; it provides a basis for investigating system perturbations, predicting experimental outcomes, and identifying a system’s most critical processes. Given the increasingly tight coupling between computational biology and biomedical research, the research community benefits when models can be easily repurposed across research groups. However, modelers use a variety of modeling languages and simulation platforms, not all of which interoperate coherently. Researchers aiming to repurpose a published model for their investigations must often resort to hand-coding the model anew, which is a time consuming and error-prone process. Although much progress has been made in developing standardized model exchange formats such as SBML^1^ and CellML^2^, fundamental roadblocks associated with model reuse remain^3–6^. Here, we identify several critical bottlenecks that impede the reuse of models and model components as well as their integration. We argue that standardizing and expanding the semantic annotations on models removes these bottlenecks by helping researchers more quickly locate models, automate model compositions, translate between modeling formats, and integrate the biological knowledge encoded in models.

Our perspective on semantic annotation is shaped by our participation in the COmputational Modeling in BIology NEtwork (COMBINE), a community of researchers developing standards for modeling in computational biology^7^. Collectively, we represent teams developing SBML, CellML, the Simulation Experiment Description Markup Language (SED-ML)^8^, the Systems Biology Graphical Notation (SBGN)^9^, the Synthetic Biology Open Language (SBOL)^10^, NeuroML^11,12^, rule-based modeling languages^13,14^, MultiCellDS^15^, the COMBINE archive^16^, the Semantic Simulation (SemSim) architecture^17^, FAIRDOMHub^18^, the SABIO-RK database^19^, as well as BioModels^20^ and the Physiome Model Repository^21,22^. Here, we present a set of agreed-upon recommendations for harmonizing semantic annotations for computational models across these development efforts. Adopting these recommendations within our communities, together with the implementation of software libraries and tools that adhere to the recommendations, will vastly extend the possibilities for interpreting, comparing, and evaluating computational models and thus enhance model reuse and repurposing.

### Semantic annotations and their utility

In the context of biological modeling, a semantic annotation is a metadata item that formally captures, entirely or in part, the meaning of a model, model component, or data element. For example, an annotation on a model variable might indicate that it represents the *concentration of cytosolic glucose in a pancreatic beta cell*, or it might describe a purely computational feature such as the *simulation time step*. To further clarify, we make a distinction between semantic annotations and other types of metadata such as curatorial annotations that describe, for example, model authorship, licensing, or provenance information. Broadly, semantic annotations are a critical feature of the vision of the Semantic Web, wherein documents are linked to metadata describing the document’s contents, thus facilitating search and retrieval as well as data interoperability^23^. Semantic annotations are used in many fields besides biological modeling, including the geosciences^24^, music retrieval^25^, and business process modeling^26^.

Semantic annotations describe the meaning of a model’s contents in a machine-accessible manner. These annotations are needed because modelers and tool developers do not use a standardized set of names to indicate the meaning of model elements. One modeler may use a variable named “X” to represent *cytosolic glucose concentration in a pancreatic beta cell*, and another modeler may use “X” to indicate *blood flow rate through the aortic valve*. In the absence of community-wide naming conventions, machine-accessible metadata is used to capture the meaning of model elements. This metadata links model elements to terms from controlled knowledge resources, allowing different software tools to recognize when two models represent the same or similar biological features, which in turn enables researchers to better align, reuse or merge models.

Annotations have contributed to the successful reuse and exploration of models and data in tasks such as comparison^27,28^, interpretation^29^, retrieval^30–32^, integration^33–37^, simulation^38^, translation between formats^29,37,39–42^ (see also <u>http://sbfc.sourceforge.net/mediawiki/index.php/SBML2BioPAX</u>), and visualization^37,43–45^. Semantic annotations are also a key component for model-driven design of synthetic biological systems where they are used in model composition tasks when constructing optimum biological systems built from models^41,45–48.^

### Challenges in model reuse and integration

Several barriers impede the reuse of models and model components. Researchers cannot easily search across multiple repositories to find models of interest, and when a researcher does retrieve a relevant model, they must determine whether it (or part of it) can be repurposed for use in their modeling work. Similarity measures and pattern-matching algorithms can help identify relevant modeling components^28,49^, but existing proposals for cross-repository search and retrieval^32^ are mostly theoretical and not yet applicable in practice.

The amount of time required to select a model for reuse increases when the researcher must choose from a large set of potentially usable models. With only limited means to compare models automatically, researchers must manually assess the content, scope, underlying assumptions of each candidate model as well as the biological questions each was designed to answer. While semantic annotations cannot yet capture the purpose for which a model was built or modified, they can make the biological content of a model explicit, and therefore help researchers decide whether or not to repurpose it. Semantic annotations can also be used to quantify the similarity between model elements^27,28,50–54^, helping to ensure that the user is presented with the most relevant models following a search. Other types of annotations, such as those that capture provenance information^55–58^, can also help researchers determine which models best meet their research needs and make comparisons between different model versions^59^.

Another set of barriers are associated with model-to-model integration. Composing new models from existing models remains a largely manual and error-prone process, and recent work has demonstrated how it can be accelerated using semantic annotations^31,34,36,60^. By examining annotations, software tools can recognize where models overlap in their biological content and then provide recommendations about how the models could be coupled.

An additional impediment to model reuse is the lack of integration between model repositories and experimental data stores. If such integration existed, modelers could readily find data appropriate for validating or constraining the models they repurpose, and experimentalists could find models for use in data analysis pipelines. If datasets and models were annotated using a standard protocol, software tools could discover relevant connections between them and accelerate research. Efforts such as the SourceData project^61^, which aims to make biological data discoverable through the use of metadata annotations, can help achieve this vision. Complementary efforts that link models with associated datasets also contribute to this goal. For example, the Simulation Database^62^ component of the JWS Online Model Repository^63^ or tellurium-web^64^ provide standardized descriptions of reproducible simulation experiments (encoded in SED-ML), manually linked to model code and, if available, original experimental datasets. Using such standardized descriptions, researchers can readily reproduce and evaluate curated simulations against experimental datasets. The opportunities for integrating models and data could greatly increase if model and data repositories such as these used a standard annotation protocol.

## Examples of current semantic annotation practices

Participants across several modeling initiatives recognize the importance of semantic annotations; however, annotation practices vary across these efforts and are in need of standardization. To illustrate, we profile the semantic annotation practices of three initiatives within COMBINE.

### BioModels

BioModels is a repository of publicly available models maintained at the European Molecular Biology Laboratory’s European Bioinformatics Institute (EMBL-EBI). The most recent BioModels release (#31) includes 640 curated models. The BioModels team has created guidelines for annotating the semantics of these models in accordance with the Minimum Information Requested In the Annotation of Models (MIRIAM)^65^ guidelines (<u>http://www.ebi.ac.uk/biomodels-main/annotationtips</u>). Annotations within curated SBML models also follow the instructions given in SBML specification documents^66^. They are primarily encoded as Resource Description Framework (RDF) statements (<u>https://www.w3.org/RDF/</u>), although the SBML format also contains explicit constructs for associating Systems Biology Ontology^38^ terms with model elements outside of the RDF content. The BioModels team has developed an in-house annotation tool for capturing the meaning of the biological aspects of SBML models using terms from a defined list of established, controlled vocabularies such as Chemical Entities of Biological Interest (ChEBI)^67^, Gene Ontology^68^ and UniProt^69^.

Furthermore, the EMBL-EBI maintains a service at <u>http://identifiers.org</u> for resolving the Uniform Resource Identifiers (URIs) used in annotations^70^. Formatted according to the identifiers.org guidelines, URIs from various biomedical knowledge resources can be resolved online. Together with the COMBINE-maintained BioModels.net “qualifiers” (<u>http://co.mbine.org/standards/qualifiers</u>), also known as “predicates” or “relations”, they allow the construction of complete semantic annotations^65^ linking elements of COMBINE formats (e.g. SBML, SED-ML or COMBINE archive metadata) to knowledge resource terms in order to define an element’s biological meaning. Figure 1 shows an RDF-based semantic annotation on an example SBML model from BioModels.

**Figure 1.**
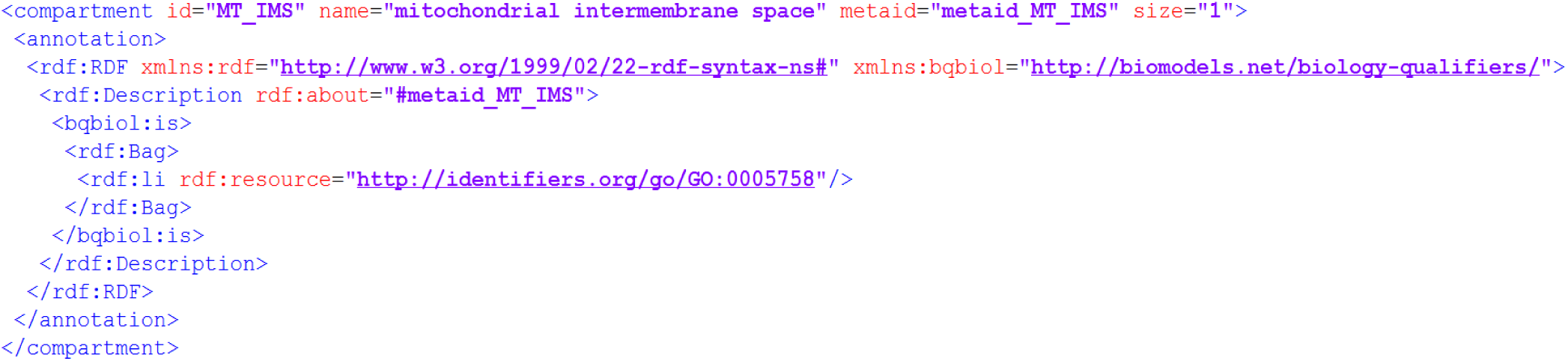
Example RDF-based annotation from SBML model BIOMD0000000239^71^ in BioModels. The *<annotation>* element defines the biological meaning of a physical compartment in the model. The RDF block within the annotation element links the compartment’s metadata identifier “metaid_MT_IMS” to the Gene Ontology term GO:0005758, which represents the mitochondrial intermembrane space. The use of the *bqbiol:is* predicate in this link indicates that the compartment is defined as the mitochondrial intermembrane space.

### The Physiome Model Repository

The Auckland Bioengineering Institute at the University of Auckland manages the Physiome Model Repository, which currently contains over 800 CellML models as well as models and simulation protocols encoded in various other formats. Annotation of CellML models is currently limited, but a collection of metadata specifications that provide recommendations and best practices for annotating models exists^72,73^. Although the CellML metadata specification states that semantic annotations should be serialized externally, current tools used in the CellML community such as OpenCOR embed RDF/XML annotations in the CellML documents themselves, using identifiers.org URI formatting and BioModels.net qualifiers.

The CellML format focuses on representing the mathematical aspects of models, and it does not include biological constructs as in SBML models. Consequently, a model’s variables must be linked to knowledge resource terms to capture the precise meaning of what a CellML model simulates. This presents challenges, as these variables represent concepts that can be very fine-grained (e.g. *concentration of cytosolic glucose in a pancreatic beta cell*), and publicly available knowledge resources do not provide adequate coverage for such concepts. Thus, the CellML group has been collaborating with the creators of the SemSim architecture, described next, to develop an annotation approach that uses *composite* annotations^17^ to describe such fine-grained concepts.

### The SemSim architecture and SemGen

The SemSim architecture is a logical framework for capturing the biophysical meaning of what is represented in a biological model. Central to this architecture are composite annotations^17^: logical statements that link multiple knowledge resource terms to precisely define a model element. The primary motivation behind the composite annotation approach is that biological models often simulate concepts that are not represented among the set of publicly available biological knowledge resources; therefore, annotators often cannot define a model element via a reference to a single controlled vocabulary term. With composite annotations, annotators can instead build a definition from multiple, more fundamental terms that are available in knowledge resources. For example, a model variable might simulate the *concentration of cytosolic glucose in a pancreatic beta cell*, but this concept is not explicitly represented in any single publicly available knowledge resource. Applying the SemSim architecture, an annotator can create a composite annotation that captures the variable’s biological meaning by linking the terms *Chemical concentration* from the Ontology of Physics for Biology (OPB)^74,75^, *glucose* from ChEBI, *Cytoplasm* from the Foundational Model of Anatomy (FMA)^76^, and *Type B cell of pancreatic islet* from the FMA.

The SemSim development group has created SemGen (<u>https://github.com/SemBioProcess/SemGen</u>), a software tool for applying SemSim-compliant composite annotations to computational models. SemGen also provides capabilities for semantics-based model composition wherein a model’s annotations are leveraged to automate merging and extraction tasks^33,34^. Using SemGen, composite annotations applied to a computational model can be stored within SemSim models, which are encoded in the RDF-based Web Ontology Language (OWL - <u>https://www.w3.org/OWL/</u>), or within SBML and CellML models as standard RDF. The SemSim development group maintains an informal protocol for annotating a model with SemGen in accordance with the SemSim framework (<u>https://github.com/SemBioProcess/SemGen/wiki/Annotation-protocol</u>). Among its guidelines are recommendations for which ontologies to use for different components of a composite annotation. For example, the OPB is recommended for physical property terms (*chemical concentration*, *fluid volume*, etc.), and the FMA for macroscopic physical entities.

As evidenced by these three initiatives, there is currently no consensus approach to semantic annotation being applied to the different projects under COMBINE. Every COMBINE initiative either uses a protocol specific to that initiative, or has no established protocol for semantic annotation. A consensus approach is essential not only for integration across COMBINE standards but also for integration with external biological data representation efforts. For example, the *open* EHR (http://www.openehr.org/) and HL7 Fast Healthcare Interoperability Resources (FHIR - https://www.hl7.org/fhir/) initiatives have established protocols for the semantic annotation of clinical data (called terminology binding). As discussed above, a consensus protocol for annotating models and data would greatly benefit researchers who apply modeling to understand clinical data. Working toward this goal, members of the *open*EHR and COMBINE communities have recently begun collaborating to coordinate their semantic annotation approaches. Thus, a primary challenge for the COMBINE community is to develop an approach to semantic annotation that will help ensure annotations among COMBINE standards are widely interoperable and consistent.

## Recommendations

Through discussions and face-to-face meetings at COMBINE meetings, we have developed the following seven recommendations for harmonizing semantic annotations:

### 1) Encode annotations as RDF; use identifiers.org URI formatting and BioModels.net qualifiers

RDF, a World Wide Web Consortium-recommended standard for representing information on the Web, has emerged as the *de facto* standard for encoding semantic annotations among the COMBINE community; all COMBINE standards currently use it. RDF has the expressivity to represent complex semantic annotations, including composite annotations, and it is a commonly-used format for storing semantic metadata in documents. RDF statements are built using subject-predicate-object triples that, as shown in Figure 1, can be used to assert relationships between model components and terms from online knowledge resources. A primer on RDF is available (<u>https://www.w3.org/TR/rdf11-concepts/</u>), and links to examples of RDF-encoded model annotations can be found in the “Example Semantic Annotations” section below.

We recommend using the identifiers.org URI format when referencing knowledge resource terms in RDF statements because identifiers.org supports a vast set of biological knowledge resources used for semantic annotation, including databases as well as ontologies, and because identifiers.org-formatted URIs are web-resolvable. We also recommend that the community uses the BioModels.net qualifiers in annotation statements that define model elements. No other standardized, centralized set of predicates exists, and the existing qualifiers provide a basic level of coverage needed for articulating semantic annotations in models.

### 2) Store annotations in a separate file

Current practice among the COMBINE community is to store semantic annotations within the same file that specifies a model’s computational aspects. This is in line with the initial MIRIAM guidelines put forth for annotating models^65^; however, there are several reasons why we recommend storing semantic annotations in a separate file. First, we wish to normalize the format in which annotations are stored across the different COMBINE standards. Currently, the exact format used to store annotations within model files differs slightly from standard to standard. Normalizing the format will simplify the development of software that provides programmatic manipulation of semantic annotations and will allow for better separation between modeling and annotation tasks. This will also remove the burden of supporting semantic annotation from the software teams that are developing software libraries for specific COMBINE standards. We recommend establishing a separate development group that will be responsible for creating and maintaining a common data model and annotation library (see Recommendation 3). If the community decides to change the way semantic annotations are stored or processed, this should not require each team of developers to adjust their software. Instead, there should be one development project focused on the programmatic manipulation of semantic annotations, and the development team for that annotation library should implement the community’s decisions.

We also recognize that different research groups may have different preferences for which knowledge resources to use for annotation. This is another important reason why we advocate storing annotations separately: externalizing annotations in a separate file allows a single model file to be referenced by multiple annotation files, allowing different research groups to describe the same modeling resource in different ways. This follows the vision of the COMBINE archive, wherein multiple types of modeling files are archived together (as in a zip file) to make simulation experiments readily reproducible and shareable among research groups^16^. When sharing models, we recommend that annotations be distributed along with the files they annotate, and COMBINE archives provide a standardized way to bundle such files together.

We recognize that storing annotations in a separate file requires keeping them synchronized. For example, if a variable identifier changes in the model file, that change should be reflected in the annotation file(s) as well. However, ensuring synchrony is an issue regardless of whether or not annotations are stored in a separate file, and we recommend that the community encourage the development of tools that help ensure coordination between a model’s computational aspects and its semantic annotations.

An additional advantage of storing annotations in a separate file is that the RDF content can be serialized in various formats including XML or Turtle, whereas currently the serialization is dictated by the model format.

### 3) Establish a dedicated group for developing a software library that supports semantic annotation standards

Programmatically creating, applying, and managing semantic annotations on models requires substantial software development. The modeling community, especially model curators, would benefit from the creation of a single dedicated group that focuses on creating and disseminating semantic annotation standards and developing standards-compliant software. The group’s role would be to promote consistency in annotation practices and help ensure that advancements in annotation standards are coordinated across the modeling community. It would also work to ensure the longevity of annotation tools beyond individual software projects within COMBINE. Given that several tools already provide model annotation capabilities, including OpenCOR^77^, SemGen^17,34^, COPASI^78^, JWS Online, CellDesigner^79^, and the SBML model annotation software used internally by BioModels, the working group’s focus would not be to create a one-size-fits-all semantic annotation toolset, but rather provide guidance and core software infrastructure to help ensure that users and developers adhere to the community’s annotation standards.

This centralized semantic annotation software package should harmonize with other annotation standards, such as the curatorial metadata standards of MIRIAM, and should support compliance-checking. We would also encourage package developers to provide multilingual support so that the lexical content associated with models and annotations can be readily converted into a wide variety of languages. This would further enhance researchers’ abilities to comprehend the meaning of model elements.

### 4) Document which knowledge resources should be used for annotation and why

The COMBINE community currently uses terms from a variety of biomedical knowledge resources to capture the meaning of model elements. Some of these knowledge resources overlap in content, and there is no broad consensus about which resources should be used for an annotation. Therefore, the same biophysical concept represented in two different models might be annotated against different knowledge resource terms. This undermines the community’s ability to compare and compose models in an automated fashion, as well as convert between standard formats. Ideally, the synonymous content of the models should be annotated using the same set of reference terms and qualifiers. Otherwise, the developers of model comparison and composition tools must rely on mappings between knowledge resources to identify synonymous terms. This is burdensome, given the increased computational cost and the challenges associated with ontology mapping and alignment^80^. It is in the interest of the community to agree upon a set of core knowledge resources that will be used for annotation, and also define the scope of use for each resource. That being said, we recognize that different research groups have different requirements for disambiguating model content. For example, UniProt might be a more useful resource compared to the Protein Ontology for a group that needs to disambiguate proteins by amino acid sequence. Furthermore, the value of different knowledge resources may change over time. Therefore, we do not propose a short list of recommended knowledge resources here. Instead, we encourage research groups to make recommendations on a group-by-group basis and provide publicly available documentation describing how specific knowledge resources should be used in annotations. This will help those outside the research group determine how to map annotations between research efforts. We also stress the importance of creating and maintaining formal mappings between knowledge resources, such as those provided by BioPortal (<u>https://bioportal.bioontology.org/</u>) and the EMBL-EBI Ontology Xref Service (<u>http://www.ebi.ac.uk/spot/oxo/</u>).

We recommend that research groups use knowledge resources that are publicly available, programmatically accessible via Web services, and mapped to other resources used within the community that overlap in content.

### 5) Establish a repository of reusable annotations

The process of annotating a model is time-consuming and susceptible to inter-annotator variability. We therefore recommend the creation of an online repository of curated, reusable annotations, and integration between this repository and software tools. The repository would allow annotators to quickly find and reuse valid, standards-compliant annotations previously applied to curated models, reducing annotation time and promoting consistency among annotators. This is especially important for applying composite annotations; they take longer to create compared to singular annotations and require linking multiple knowledge resource terms, resulting in more opportunities for inter-annotator inconsistency. An additional benefit of such a repository is that the annotations it contains could be used for annotating experimental datasets. We envision this repository would mine RDF annotation statements from curated models (e.g. in BioModels or the PMR) and integrate with annotation software tools so that annotators can quickly link elements in their models/datasets to knowledge resource terms by reusing these statements.

Such a repository could also provide the basis for automating the annotation process; once an annotator begins annotating a model, software tools could search the repository for curated models that have the same annotations and make suggestions for unannotated model elements. This approach would also improve inter-annotator consistency; see Recommendation 4, above.

By utilizing the mappings that exist between knowledge resources (e.g., between ChEBI and the Kyoto Encyclopedia of Genes and Genomes^81^), a centralized repository of annotations could also identify and auto-generate synonymous annotations. This would help minimize redundant annotations within the repository and allow annotators to retrieve annotations that only use references to their preferred knowledge resources.

As discussed in Recommendation 3, we encourage the developers of an annotation repository to provide multilingual support.

### 6) Ensure high-quality semantic annotations through training and quality control processes

The success of new software tools for expediting cross-repository model retrieval and model composition depends on linking models to thorough, precise semantic descriptions. Therefore, we recommend that annotators, whether they are the modelers themselves or curation team members, receive training so that their annotations are specific, complete and as consistent as possible across repositories. Annotators must not only thoroughly understand the meaning of a model’s computational elements, but also the scope, organization, and limitations of the knowledge resources used for semantic annotation as well as the technical aspects of encoding annotations.

We also encourage all groups annotating models to develop quality control protocols that ensure annotations are encoded correctly, to identify overly generic, overly specific, and missing annotations, and to check that the annotations accurately capture the meaning of the model elements to which they are linked. We also recommend the enhancement of existing annotation software tools^36,82^ so they prevent the application of inconsistent annotations and can auto-suggest annotations. Another interesting approach for promoting high-quality annotations is the use of Web 2.0 advancements and collaborative technologies to evaluate semantic annotations with user-friendly and interactive infrastructures (see, for example, Kalbasi et al.^24^).

### 7) Establish and maintain collaborations with knowledge resource developers

Modelers do not always organize a system for study in the same way as knowledge resource developers, and sometimes the biological concepts represented in a model are not present in existing knowledge resources. Many biological concepts in models are too fine-grained to be included and maintained in knowledge resources aiming to provide broader classifications for biophysical phenomena. Thus, annotating a model may require sending term requests to resource developers. Therefore, the modeling community should establish and maintain connections with these developers so that terms needed for annotation can be added to resources or modified in a timely fashion. For example, the Kinetic Simulation Algorithm Ontology (KiSAO)^38^ term submission system on the COMBINE website (<u>http://co.mbine.org/standards/kisao</u>) connects to a SourceForge ticket system. An ongoing dialog between modelers and resource developers will also help modelers better understand the scope and content of knowledge resources, which is a critical component of Recommendation 6.

## Example Semantic Annotations

To provide concrete examples of semantic annotations that adhere to our recommendations, we have created two example COMBINE archives that contain annotated models. These archives are included as Supplementary Information, and their latest versions are available at the COMBINE GitHub repository (<u>https://github.com/combine-org/Annotations</u>). One of these archives contains an SBML model retrieved from BioModels (BIOMD0000000176) that simulates glycolysis in yeast^83^. The other archive contains a CellML model retrieved from the Physiome Model Repository that simulates hemodynamics in the human circulatory system^84^. These two models were chosen because, together, they provide examples of semantic annotations across physical scales, and because the phenomena they simulate will be familiar to many biological researchers. In the future, we plan to create example semantic annotations for additional COMBINE standards (e.g. NeuroML and SBGN) and post them to the COMBINE GitHub repository as well.

We have serialized semantic annotations for both of our example models as separate files within COMBINE archives. The annotations within these standalone files are encoded as RDF subject-predicate-object statements and the URIs for the subjects in these statements serve as pointers to the specific model elements that are annotated. To construct these pointers, we use the relative model file name for the URI base and the “metaid” attribute on the annotated model element for the URI fragment. For example, the URI referencing the cytosol compartment in the SBML model is ./BIOMD0000000176.xml#_525523. This approach allows us to uniquely reference model elements within a COMBINE archive, even if elements from two different models share the same metaid. These URI pointers establish the link between the model file and its annotations, and the original model code is left unedited. By asserting links this way, a model file does not have to be updated when it is annotated, and synchronization of pointers across multiple files is not required.

Our examples are based on initial progress towards a full technical specification of how external annotation files should be implemented for COMBINE standards. However, this specification is still under development and a number of implementation details must be resolved before it can be completed, including how annotations should be implemented in BioPAX^85^ and SBOL documents. Regardless, we hope that these examples will provide an initial reference point for standardization of a COMBINE-wide semantic annotation protocol. We invite those interested in participating in this standardization process to join the “COMBINE-annot” online forum (<u>https://groups.google.com/forum/#!forum/combine-annot</u>), which provides a venue for addressing issues related to the annotation of models and data.

## Discussion

As discussed above, semantic annotations can accelerate model reuse by enhancing search and retrieval of models. If a standard semantic annotation protocol is applied to publicly available models, the modeling community can develop more advanced tools that can search across model repositories, improve the relevance of search results, and provide users with information that helps them decide whether an existing model is suitable for reuse in a modeling project. A common annotation protocol is also a critical component of model composition. Applying the protocol, model-merging tools such as SemGen can recognize the biological commonalities between models and then use that information to compare the models’ biological content and guide their assembly. Without a harmonized approach to semantic annotation, modelers would be required to manually identify the biological overlap between models.

Given that data is an essential component of model-based research, we strongly encourage the computational biology community to develop protocols for data annotation. We hope that the annotation approach articulated here, and implemented in the example annotation files described above, provides a basis from which to build such protocols. It is in the interest of both modelers and experimentalists to adopt a shared approach for semantic annotation: by linking annotations in models and data sources, modelers could discover valuable datasets for tuning and validating their models, and experimentalists could discover simulation models that integrate knowledge about their field of study into systems-level perspectives.

We recognize that it takes time and effort to annotate models accurately and consistently. Therefore, we urge the community to develop incentives and tools to make it easier and faster to annotate models. Incentives might include giving credit to annotators in model metadata and on model repositories. New tools that automate compliance-checking of annotations (e.g. <u>https://normsys.h-its.org/validate</u>) could help flag errors, omissions, and overgeneralizations.

Despite these challenges, we believe that semantic annotation is one of the keys for transforming the entire field of biological modeling into one in which models do not languish within single research labs, but are readily shared among the broader research community, easily incorporated into new modeling projects, and contain reusable components. As the biotechnology community undertakes more investigations into systems-level biology, standardized semantic annotations will reduce the activation energy required to initiate, repurpose, and extend model-based research projects. We hope that the recommendations articulated here help move the greater modeling community, including groups outside of COMBINE, towards the goal of broad model reuse and interoperability.

## Acknowledgements

MLN, DLC and JHG were supported by the United States of America’s National Institutes of Health (NIH) grant R01 LM011969. MLN was also supported by NIH grant P41 GM109824. DW was supported by the German Federal Ministry of Education and Research (BMBF) (SEMS, 031 6194). MK was supported by BMBF (LiSyM, 031L0054). DN was supported by an Aotearoa Foundation Fellowship. RK and KA were supported by the Medical Technologies Centre of Research Excellence (MedTech CoRE). AD, MH and NL were supported by the National Institute of General Medical Sciences (NIH/NIGMS) grant R01 GM070923. SC was supported by NIH grants R01 MH106674 and R01 EB021711. MdA was supported by the European Commission’s (EC) grant 731001. MG was supported by the German Federal Ministry for Economic Affairs and Energy (BMWi) (NormSys, 01FS14019), by BMBF (LiSyM, 031L0056), and by the Klaus Tschira Foundation (KTS). CM was supported by the United States of America’s National Science Foundation under grants 1522074, CCF-1218095, and DBI-1356041. HMS was supported by NIH grant R01 GM123032. JLS was supported by the Department of Science and Technology/National Research Foundation (DST/NRF) in South Africa (NRF-SARCHI-82813) and by the Biotechnology and Biological Sciences Research Council in the United Kingdom (BBG0102181, BB/I004637/1, BB/M013189/1). VT was supported by the Norwegian University of Science and Technology’s Strategic Research Area "NTNU Helse." Any opinions, findings, and conclusions or recommendations expressed in this material are those of the author(s) and do not necessarily reflect the views of the funding agencies.

## Author Contributions

MN and DW wrote manuscript drafts and prepared the final submission. All authors contributed content and/or edits to manuscript drafts. All authors participated in outlining and refining the recommendations articulated in the manuscript. MN and MK prepared the example semantic annotation files.

